# Physiological Reconstitution of Microtubule Doublets

**DOI:** 10.1101/2025.08.03.668368

**Authors:** Ming Li, Guanghan Chen, Zhe Chen, Zhengyang Guo, Zi Wang, Yongping Chai, Wei Li, Guangshuo Ou

## Abstract

The formation of microtubule doublets (MTDs) is a foundational step in cilia biogenesis, yet how B-tubule nucleation is initiated at the molecular level remains elusive. Here, we identify FAP53 as a crucial factor that promotes B-tubule assembly. Molecular dynamics simulations revealed that FAP53 alleviates steric hindrance from the α-tubulin C-terminal tail and stabilizes B-tubule docking at the A-tubule surface. In vitro reconstitution demonstrated that recombinant FAP53 is sufficient to drive MTD formation under physiological tubulin conditions. In cultured HeLa cells, co-expression of CFAP53 and CFAP20—an inner junction protein—induced ectopic MTD-like structures in the cytoplasm. Furthermore, a structurally homologous protein in *C. elegans*, WFAP-53, localized to sensory cilia; however, when overexpressed, it triggered ectopic MTD formation in neuronal dendrites and concomitantly led to sensory cilia disassembly. These findings uncover a conserved mechanism of B-tubule initiation and underscore the necessity for spatially restricted expression of MTD assembly factors during ciliogenesis.

## Introduction

Cilia are conserved organelles essential for motility, signaling, and tissue homeostasis ^1–5^. Their structural core, the axoneme, is composed of microtubule doublets (MTDs), with each doublet consisting of a complete 13-protofilament A-tubule and an incomplete 10-protofilament B-tubule ^1–5^. MTDs form the basis of both motile (“9+2”) and primary (“9+0”) cilia, serving as platforms for dynein-driven motility or as scaffolds for signaling and cargo transport ^1–5^. The unique, asymmetric architecture of MTDs is central to ciliary function, but the molecular logic of their assembly remains poorly understood.

Reconstituting MTDs in vitro using native components is a powerful approach to dissect the design principles underlying the construction of complex cellular structures. It also has direct clinical relevance: loss of MTDs disrupts cilia, and reassembly begins with doublet formation during cilium regeneration ^6–8^. However, tubulin heterodimers preferentially form singlet microtubules, and reconstituting physiologically relevant MTDs has been challenging due to the energetic and geometric barriers specific to B-tubule nucleation ^9–11^. These challenges suggest the involvement of accessory proteins that regulate doublet-specific architecture.

One major obstacle to B-tubule assembly is the α-tubulin C-terminal tail (CTT), which creates a steric barrier that inhibits lateral attachment of the B-tubule to the A-tubule surface ^9,10^. Although enzymatic removal of tubulin CTTs facilitates the formation of doublet-like structures in vitro ^10,12^, these artificial treatments bypass native regulatory mechanisms and fail to recapitulate physiological assembly. Recent cryo-electron microscopy studies have revealed a dense network of microtubule inner proteins (MIPs) lining the lumens of both A- and B-tubules, particularly concentrated at the inner and outer junctions where tubule interactions are most critical ^13–18^. Among outer junction MIPs, FAP53 emerges as a compelling candidate for regulating B-tubule initiation. It is highly conserved across ciliated species, and its vertebrate homologs (CFAP53/CCDC11) are essential for dynein arm docking, ciliary motility, and left-right axis specification ^19–22^. Human mutations in CFAP53 cause a spectrum of ciliopathies, including situs abnormalities, infertility, and respiratory dysfunction ^20–26^; yet, the molecular function of FAP53 in doublet formation has remained poorly defined. Structurally, FAP53 spans the tripartite interface formed by protofilaments A10, A11, and B01, placing it in a prime position to overcome the CTT-imposed steric barrier and to nucleate B-tubule formation via both lateral and longitudinal contacts ^13,16^. These features make FAP53 a strong mechanistic candidate for initiating and stabilizing the outer junction during MTD biogenesis.

Here, we reconstituted MTDs in vitro using native α/β-tubulin and recombinant FAP53, establishing a minimal and physiologically relevant system that recapitulates the earliest steps of B-tubule initiation. By integrating molecular dynamics simulations, biochemical reconstitution, and in vivo validation in both human cells and *C. elegans*, we demonstrate that FAP53 relieves CTT-induced steric hindrance and stabilizes the outer junction architecture. These findings establish FAP53 as a core initiator of MTD formation and offer a modular platform for engineering asymmetric microtubule structures—advancing synthetic approaches to cytoskeletal design. Moreover, this system provides a foundation for dissecting the etiology of ciliopathies linked to CFAP53 dysfunction, and for functionally testing disease-associated mutations in a controlled, reconstituted context.

## Results

### Molecular Dynamics Simulations Reveal a Dual Role for FAP53 in B-Tubule Initiation

To gain insights into whether and how FAP53 facilitates B-tubule assembly, we constructed a structural model of the outer junction based on the high-resolution cryo-EM structure of microtubule doublets (PDB: 6U42 ^16^) (Fig. 1A). The model encompassed protofilaments A10– A11, the B01 dimer, and the N-terminal segment of FAP53 (residues 2-40). To more accurately approximate the native molecular interface, we incorporated flexible regions not resolved in the original cryo-EM map, including the α/β-tubulin CTTs and the α-tubulin H1–S2 loop. Initial Rosetta-based energy minimization revealed a steric clash between the α-tubulin CTT and the FAP53 N-terminus (Fig. 1B), echoing our previous findings that the α-tubulin CTT impedes B01 dimer docking on the A10–A11 surface and thereby hinders B-tubule nucleation ^9^. This observation led us to hypothesize that FAP53 could act to neutralize this steric hindrance and stabilize B01 engagement.

**Figure 1.**
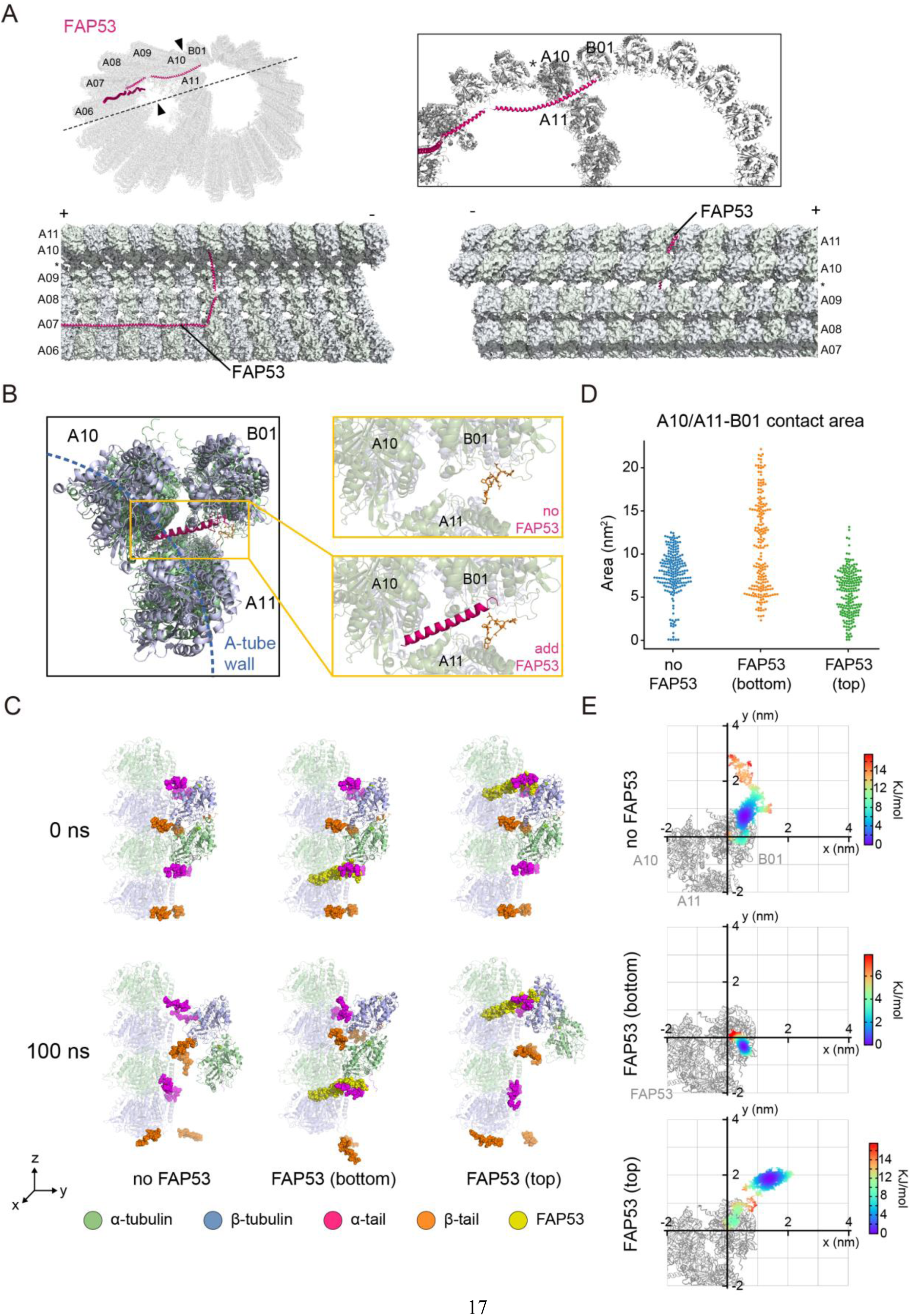
Molecular dynamic simulation predicts that FAP53 mitigates the inhibitory effect of CTT on formation of MTs assembly. (A) The structure model of MTD (adapted from PDB ID: 6U42). Front view and side view are displayed. FAP53 is highlighted as magenta. (B) Structure model of local energy minimization of FAP53. Steric clash between the α-tubulin CTT (orange) and the N-terminal of FAP53 (magenta) is showed. (C) Side views of the representative simulation result in different cases. The α-/β-tubulin CTTs and FAP53 N-terminus are represented by spheres to indicate the Van der Waals radii, while the remaining parts are depicted using a cartoon representation to highlight the secondary structure. The initial snapshot (0 ns) captures the conformation after NPT equilibrium, followed by the snapshots at the end of 100 ns simulations. (D) The contact area between B01 dimer and A10-A11 patch in different cases. The first 50 ns of the simulations were discarded as the equilibration phase. The contact area was calculated every 0.5 ns during the last 50 ns. (E) Free energy landscape inferred from the distribution of center of mass coordinate within the 100 ns simulations of each case. The probability density on the X-Y plane was estimated by kernel density estimation and the free energy was calculated by Boltzmann inversion. The origin of the coordinates is the starting position.

To test this hypothesis, we performed full-atom molecular dynamics (MD) simulations on two distinct configurations of the outer junction: one with FAP53 bound at the plus-end of the B01 heterodimer (“top” model), and the other at the minus-end (“bottom” model). Each simulation was run for 100 ns with positional restraints applied to the α-carbon atoms of A10–A11 to mimic the rigidity of the microtubule lattice (as in our previous study ^9^). Strikingly, in the absence of FAP53, the B01 dimer dissociated entirely from the A-tubule interface (Fig. 1C, E; Video S1), confirming that stable binding is energetically unfavorable under native conditions. By contrast, in both FAP53-containing models, the B01 dimer remained persistently docked throughout the simulation (Fig. 1C, E; Video S2 and S3), indicating that FAP53 plays a stabilizing role at the outer junction.

Mechanistically, the N-terminal segment of FAP53 appears to shield the α-tubulin CTT, preventing direct clashes with the incoming B01 dimer and simultaneously contributing favorable surface interactions that facilitate dimer docking. Among the two configurations, the bottom model—featuring FAP53 at the minus-end of B01—exhibited greater structural stability (Fig. 1E), consistent with enhanced compatibility between FAP53 and the B01 minus-end interface. Contact area analysis further supported this observation: the B01 dimer in the bottom model maintained more extensive and persistent interactions with A10–A11 protofilaments (Fig. 1D; Fig. S1). Together, these results suggest a dual mechanism by which FAP53 promotes B-tubule assembly: first, by sterically shielding the α-tubulin CTTs to alleviate an intrinsic barrier to nucleation, and second, by providing a structurally complementary docking interface that stabilizes the B01 dimer at the A-tubule surface. These findings implicate FAP53 as a critical co-factor in establishing the outer junction architecture of microtubule doublets.

### In Vitro Reconstitution Demonstrates that FAP53 Promotes MTD Assembly

To examine the functional role of FAP53 in B-tubule formation, we reconstituted MTDs in vitro using purified porcine brain α/β-tubulin and recombinant *Chlamydomonas* FAP53 tagged with a 6×His epitope (Fig. 2A). Building on the established MTD assembly protocol ^9,10^, we first polymerized stable singlet microtubules, then incubated them with either tubulin alone or with tubulin supplemented with recombinant FAP53. In reactions lacking FAP53, the assembly yielded primarily singlet microtubules. In contrast, inclusion of FAP53 markedly increased the formation of MTD-like structures (Fig. 2B). Cryo-electron tomography (cryo-ET) analysis revealed that these doublet structures comprised B-tubules nucleated laterally along the A-tubules— recapitulating the native outer junction architecture observed in motile cilia (Fig. 2C). Notably, a subset of reconstituted MTDs exhibited S-shaped curvature in their B-tubules, likely arising from heterogeneity in protofilament number or lateral curvature strain during B-tubule elongation.

**Figure 2.**
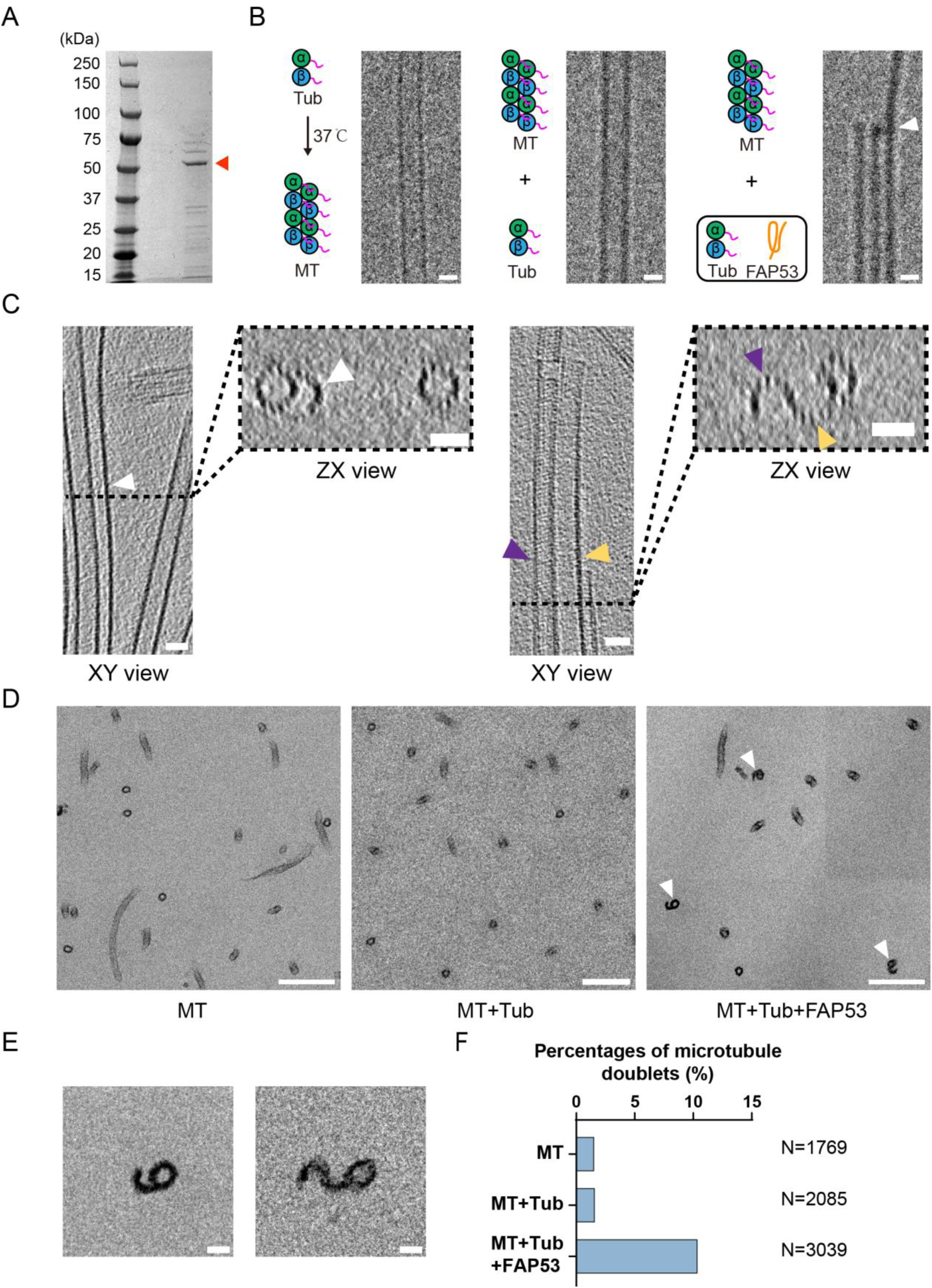
FAP53 promotes MTD formation in vitro. (A) Coomassie blue staining of purified FAP53 (indicated by red triangle). (B) Representative cryo-EM images of microtubules (MT), microtubules incubated with tubulin (MT+Tub), and microtubules incubated with tubulins and FAP53 (MT+Tub+FAP53). Arrowhead indicates MTD. Scale bars: 25 nm. (C) Representative imaged of cryo-ET sections of MT+Tub+FAP53. Arrowheads indicate MTD. Scale bars: 25 nm. (D) Representative images of cross sections of chemical-fixed microtubule polymers in MT, MT+Tub, and MT+Tub+FAP53. Arrowheads indicate MTD. Scale bars: 200 nm. (E) Representative images of MTDs in MT+Tub+FAP53. Scale bars: 25 nm. (F) Percentage of MTD formation in vitro. N indicates the total microtubules observed.

To quantitatively assess MTD formation, we performed chemical fixation followed by resin embedding and thin-section electron microscopy. In the absence of FAP53, MTDs accounted for only ∼1.6% of total microtubule structures. Upon FAP53 addition, this proportion increased significantly to ∼10% (Fig. 2D, F), confirming a stimulatory role in MTD nucleation. Consistent with our cryo-ET observations, both canonical and S-curved MTDs were present in the FAP53-supplemented reactions (Fig. 2E). These findings demonstrate that FAP53 is not only structurally poised to facilitate B-tubule initiation—as predicted by molecular modeling—but also functionally sufficient to drive MTD assembly in vitro. This reconstitution establishes FAP53 as a critical cofactor in the early stages of axonemal doublet formation, bridging structural prediction with experimental validation.

### CFAP53 and CFAP20 Cooperatively Induce Ectopic MTD Formation in the Cytoplasm of HeLa Cells

To determine whether CFAP53 can initiate B-tubule assembly in a cellular context, we ectopically expressed human CFAP53 in HeLa cells. When expressed alone, CFAP53 localized to cytoplasmic tubule-like structures that did not overlap with the endogenous microtubule network (Fig. 3A), suggesting limited intrinsic affinity for native microtubules in vivo. This observation raised the possibility that additional factors, particularly those involved in inner junction stabilization, might be required to support MTD-like architecture in vivo.

**Figure 3.**
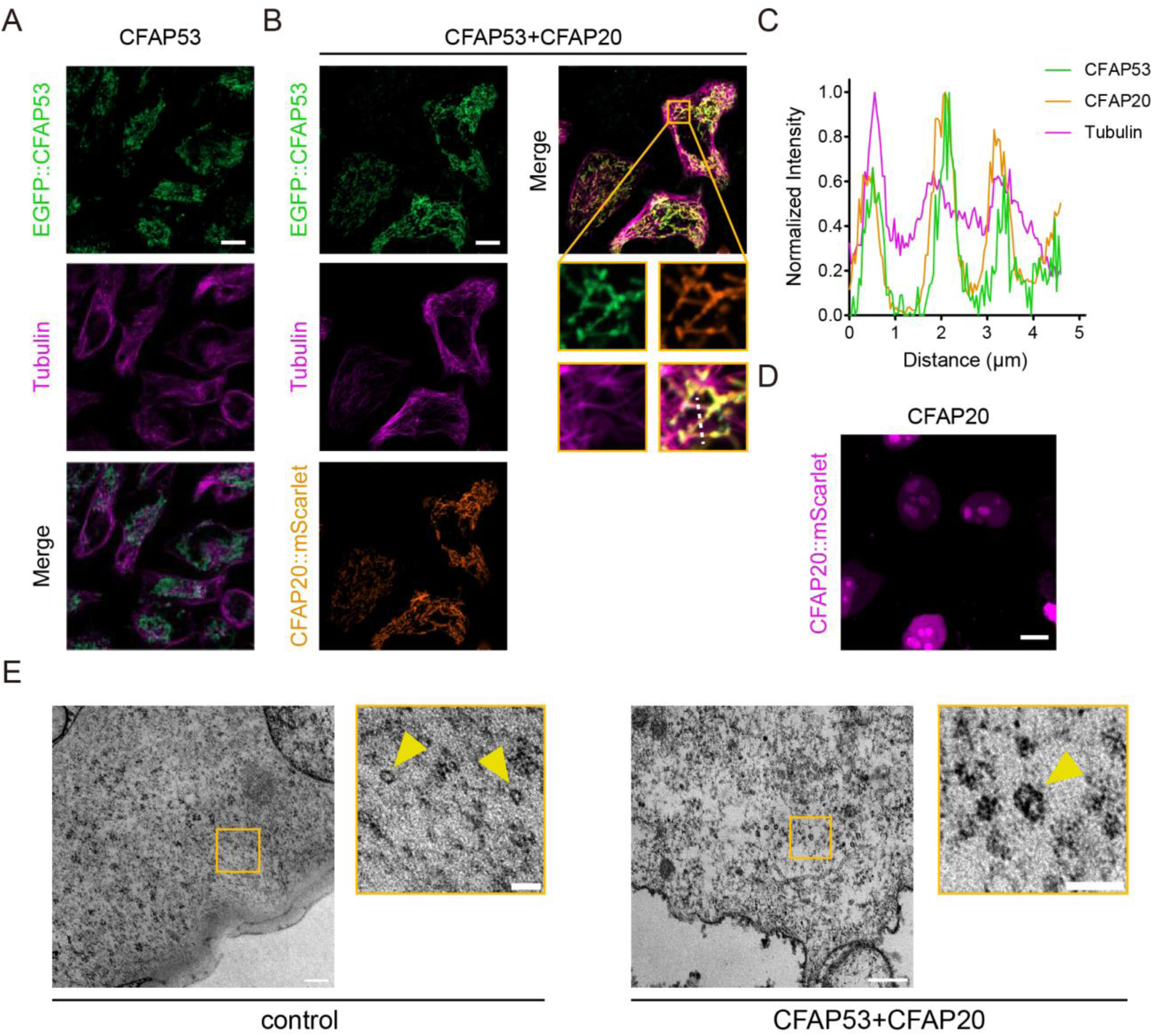
Overexpression of CFAP53 and CFAP20 induces ectopic MTDs in human cells. (A) Representative confocal images of HeLa cells expressing EGFP::CFAP53, counterstained with Tubulin Tracker Deep Red to visualize the endogenous microtubule network. Scale bar, 10 μm. (B) Representative images of HeLa cells co-expressing EGFP::CFAP53 and CFAP20::mScarlet, showing colocalization with the microtubule cytoskeleton. Scale bar, 10 μm. (C) Line-scan intensity profiles of EGFP::CFAP53 (green), CFAP20::mScarlet (orange), and Tubulin Tracker (magenta) signals across the microtubule regions (along the dashed line in panel B). All fluorescence intensity profiles are normalized to their maximum. (D) Representative image of HeLa cells expressing CFAP20::mScarlet alone, showing localization in nuclei. Scale bar, 10 μm. (E) Representative TEM images (cross sections) of control and CFAP53+CFAP20 overexpression cells. High magnification on the right side shows the microtubule singlets or MTDs (arrowheads). Scale bars: 200 nm (low magnification), 50 nm (high magnification).

Given the cooperative nature of A- and B-tubule interaction in axonemal doublets, we next tested whether CFAP20—a conserved component of the inner junction—could enhance CFAP53 recruitment and support ectopic doublet formation. Consistent with prior reports ^27^, CFAP20 expressed alone localized primarily to the nucleus and failed to associate with cytoplasmic microtubules (Fig. 3D). However, co-expression of CFAP53 and CFAP20 led to partial co-localization of both proteins along microtubule-like cytoplasmic filaments (Fig. 3B, C), indicating a synergistic interaction that promotes their recruitment to the microtubule network.

To assess whether these assemblies corresponded to bona fide MTDs, we performed transmission electron microscopy (TEM) on ultra-thin sections of transfected cells. Control cells exhibited only singlet microtubules, as expected. By contrast, a subset of cells co-expressing CFAP53 and CFAP20 displayed clear doublet-like microtubule structures, characterized by a secondary tubule laterally appended to a primary one in the cytoplasm of HeLa cells (Fig. 3E; additional examples in Fig. S2). Although these structures were observed at low frequency—likely reflecting the absence of other axonemal components or the lack of spatial constraints typically provided by the ciliary environment—their appearance demonstrates that CFAP53 and CFAP20 are together sufficient to nucleate elements of doublet architecture in the cytoplasm of non-ciliated cells. These results reveal a cooperative mechanism between CFAP53 and CFAP20 in promoting B-tubule assembly and underscore their sufficiency in initiating MTD formation outside the native ciliary compartment. This provides compelling in vivo evidence supporting their functional roles at the outer and inner junctions, respectively.

### A FAP53-Like Protein in *C. elegans* Drives Ectopic MTD Formation in Sensory Neuron Dendrites

To explore the evolutionary conservation of FAP53 function in animals, we sought to identify a structural homolog in *Caenorhabditis elegans*. Although conventional sequence homology searches failed to reveal a direct FAP53 ortholog, structural similarity screening using the ADAMS platform ^28^ identified F48E3.9 as the top candidate (Fig. 4A). Based on its predicted structural resemblance to the conserved N-terminal domain of FAP53, we named this gene *wfap-53* (worm FAP53). Single-cell RNA-sequencing data revealed that *wfap-53* expression is restricted to ciliated sensory neurons ^29^ (Fig. 4B), consistent with a potential role in cilia biogenesis. To examine its subcellular localization, we generated a translational reporter construct (*Pwfap-53::wfap-53::GFP*) and microinjected it at low concentration (5 ng/μL) into the *C. elegans* germline. The WFAP-53::GFP fusion protein localized to the base and transition zone of sensory cilia (Fig. S3A), indicating its functional incorporation into the ciliary compartment.

**Figure 4.**
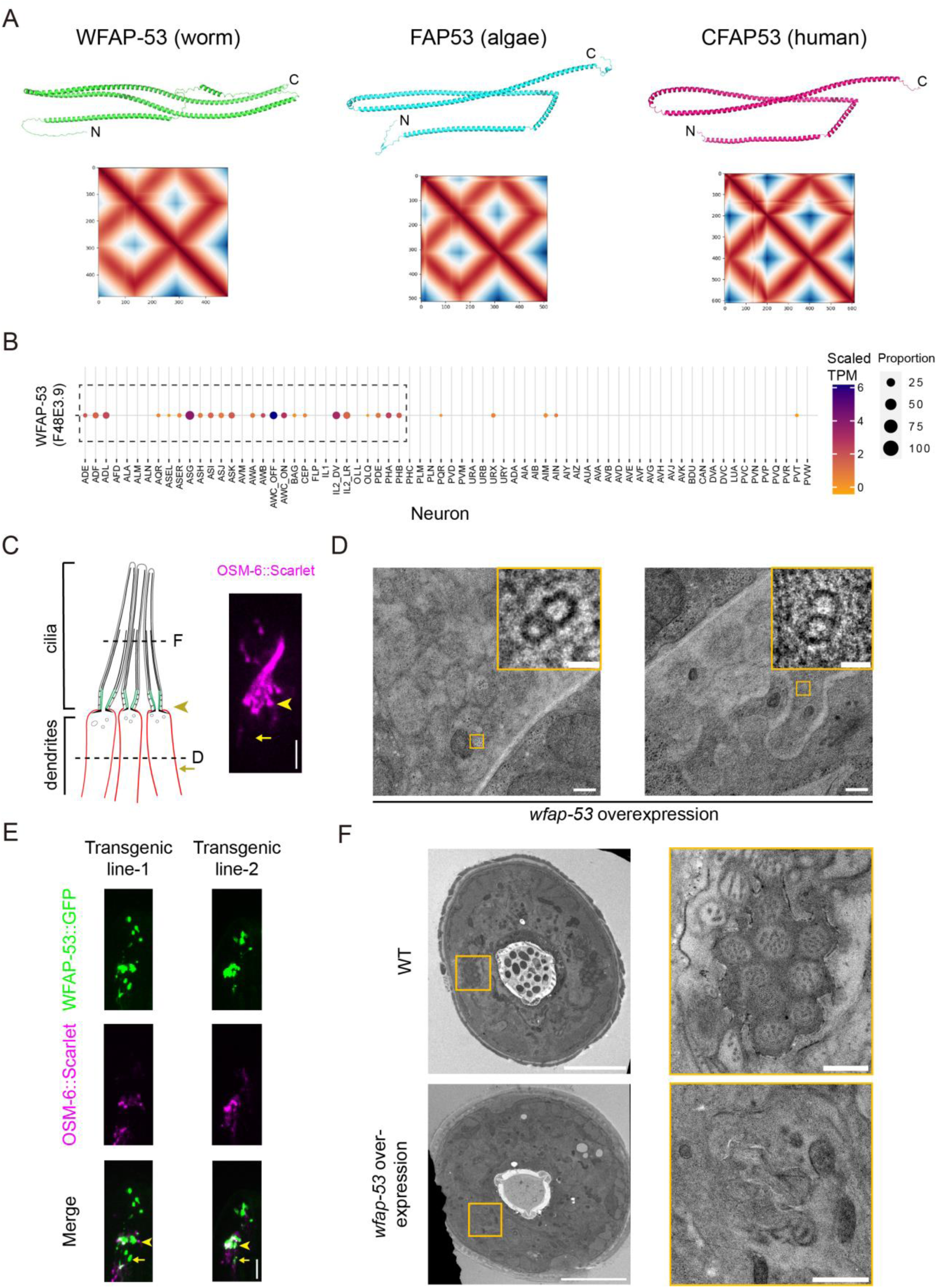
Overexpression of FAP53 structural homolog *wfap-53* induces ectopic MTDs in *C. elegans*. (A) Comparison of worm WFAP-53, Chlamydomonas FAP53 and human CFAP53 structures (predicated by AlphaFold3). The respective distance matrices (generated by ADAMS) are showed below. (B) Heatmap showing the expression levels *wfap-53* in *C. elegans* ciliated sensory neurons. The plot was generated from the CeNGEN website (https://cengen.shinyapps.io/CengenApp/). Sensory neurons are framed by dashed lines. The size of each circle corresponds to the proportion of neurons in each cluster expressing a particular gene. TPM, transcripts per million. (C) Left: schematic diagram of the amphid cilia and dendritic structure of *C. elegans*. The dashed lines indicate the location of the slices represented in Figures D and F; Right: amphid cilia in wild type *C. elegans* labeled by OSM-6::Scarlet. Arrowhead indicate the ciliary base; Arrow indicates the dendrite. Scale bar: 5 μm. (D) Representative TEM images (cross sections) of dendrites in *wfap-53* overexpression animals. High magnification on the right side shows the MTDs in dendrites. Scale bars: 200 nm (low magnification), 20 nm (high magnification). (E) Amphid cilia in *wfap-53* overexpression animals. Arrowheads indicate the ciliary base; Arrows indicate the dendrites. Scale bar: 5 μm. (F) Representative TEM images (cross sections) of amphid channel cilia in wild type and *wfap-53* overexpression animals. Scale bars: 5 μm (low magnification), 500 nm (high magnification).

To assess the effects of *wfap-53* overexpression, we increased the injection concentration to 50 ng/μL and examined amphid sensory neurons by TEM (Fig. 4C). Remarkably, we observed the formation of ectopic MTD-like structures within neuronal dendrites—compartments that do not normally support doublet assembly (Fig. 4D; additional examples in Fig. S3B). These ectopic MTDs resembled canonical A–B microtubule doublets in morphology and position, suggesting that WFAP-53 is sufficient to induce doublet-like architecture in a non-ciliary context. Notably, *wfap-53* overexpression induced severe ciliary abnormalities. Using the ciliary marker OSM-6::Scarlet, we observed truncated or missing middle and distal axonemal segments in sensory neurons (Fig. 4E). TEM analysis confirmed structural disruption of the axoneme, with deviation from the stereotypical 9+0 configuration seen in wild-type animals (Fig. 4F). Instead, MTDs were aberrantly assembled within the dendritic shaft, further supporting the notion that misregulated WFAP-53 expression redirects tubulin polymerization away from the cilium.

These findings establish *wfap-53* as a functional homolog of FAP53 in *C. elegans* and demonstrate its ability to promote MTD formation outside the canonical ciliary compartment. Moreover, the deleterious consequences of its overexpression underscore the requirement for precise spatial regulation of FAP53-like proteins during ciliogenesis. Together, these results highlight the evolutionary conservation of the FAP53 mechanism and its context-dependent impact on microtubule architecture.

## Discussion

Our study identifies FAP53 as a conserved core assembly factor that initiates MTD formation through a structurally defined and physiologically relevant mechanism. Using molecular dynamics simulations, we show that FAP53 alleviates steric hindrance imposed by the α-tubulin CTT, facilitating stable docking of the B01 protofilament onto the A-tubule surface. Unlike previous approaches relying on chemically modified or truncated tubulin, we demonstrate that recombinant FAP53 alone is sufficient to promote MTD-like assembly in vitro using full-length, native α/β-tubulin under near-physiological conditions. These results directly address a longstanding gap in our understanding of how B-tubule initiation is spatially and structurally orchestrated at the outer junction (OJ), and provide a minimalist yet tractable system for reconstructing the molecular logic of axonemal architecture.

By integrating *in silico* modeling, *in vitro* reconstitution, and *in vivo* functional validation across diverse systems, we establish a conserved mechanism by which FAP53 promotes MTD formation. FAP53 orthologs are consistently localized to the OJ across ciliated organisms—from *Chlamydomonas* to mammals—as shown by cryo-EM studies of flagella ^16^, tracheal cilia ^13^, and sperm axonemes ^15^. Our data show that *Chlamydomonas* FAP53 reconstitutes MTDs in vitro, human CFAP53 induces ectopic doublets in HeLa cells when co-expressed with CFAP20, and the *C. elegans* homolog, WFAP-53, triggers MTD assembly in dendritic compartments of sensory neurons. These findings highlight a deeply conserved strategy for building asymmetric microtubule-based structures, and demonstrate that key features of MTD nucleation can be reconstituted even outside the canonical ciliary environment.

Importantly, this work lays the foundation for mechanistically dissecting ciliopathies linked to defects in doublet assembly. CFAP53 (CCDC11) is linked to a spectrum of human disorders, including situs inversus, male infertility, and primary ciliary dyskinesia ^20–26^. While prior studies have implicated these proteins in dynein arm docking, our findings reveal their upstream role in organizing the OJ and initiating B-tubule formation. The ability to reconstruct this process using purified components opens new avenues for modeling disease-associated mutations and testing their structural and functional impact.

Although FAP53 is sufficient to trigger B-tubule initiation, the efficiency of reconstitution (∼10%) and the frequent occurrence of incomplete B-tubule closure underscore the necessity of additional cofactors for robust and complete doublet formation. Inner junction proteins such as CFAP20 and PACRG stabilize A1–B10 interactions and likely contribute to B-tubule sealing—steps not recapitulated in our minimal system ^30–33^. Additional outer junction MIPs such as FAP127 and FAP141 are predicted to play complementary roles: FAP53 binds the A6–A7 groove, FAP127 the A7–A8 interface, and FAP141 provides lateral support without longitudinal anchoring ^13,16^. This spatial partitioning suggests a sequential assembly logic in which FAP53 initiates B-tubule formation, and other MIPs stabilize elongation and closure. This model is supported by genetic studies in vertebrates, where loss of individual MIPs produces mild defects, while compound mutants exhibit severe axonemal disorganization and near-complete loss of outer dynein arms ^20^. Additional B-tubule luminal proteins such as FAP45, FAP52, and FAP77 also contribute to MTD stability. Their absence results in detachment of the B-tubule from the A-tubule wall or collapse of the OJ interface ^14,34^. Incorporating these components into future reconstitution assays will be crucial for reconstructing a fully functional MTD and for testing how specific modules contribute to mechanical robustness and ciliary beat regulation.

At the structural level, our findings suggest that FAP53 undergoes conformational remodeling upon engagement with the A–B tubule interface. While AlphaFold predicts a flexible, four-helix bundle in isolation, cryo-EM maps of native axonemes consistently show FAP53 in an L-shaped conformation, with the C-terminal domain inserted into the A6–A7 groove and the N-terminal helix spanning the OJ to engage B01 ^13,16^. This discrepancy points to regulated folding during incorporation, possibly mediated by chaperones or post-translational modifications. Elucidating the biogenesis of FAP53 itself—and how it is targeted to and stabilized at the OJ—will be important for understanding how axonemal components achieve their mature conformation and spatial positioning.

Future studies using proximity-labeling proteomics and structure-guided mutagenesis can further define the interactome and folding landscape of FAP53, and identify additional regulators that ensure fidelity during MTD assembly. More broadly, our study establishes a framework for reconstituting asymmetric microtubule-based architectures from native components, providing a template for understanding how molecular asymmetry is engineered in other organelles. As cilium dysfunction lies at the root of numerous human disorders, including polycystic kidney disease, retinal degeneration, and hydrocephalus ^35,36^, these insights carry implications beyond basic cell biology—offering new experimental systems and therapeutic entry points for targeting ciliopathies at their structural core.

## Materials and Methods

### *C. elegans* strain culture

*C. elegans* strains were cultured on nematode growth medium (NGM) plates seeded with *Escherichia coli* strain OP50 at 20 °C. All animal experiments were conducted in compliance with governmental and institutional guidelines. A summary of all strains and plasmids used in this study is provided in Table S1.

### Strain Construction

The *wfap-53* overexpression construct *(Pwfap-53::wfap-53::GFP)* was engineered through homologous recombination using the In-Fusion HD Cloning System (Clontech, 639648). Specifically, a 1309 bp genomic fragment containing the native *wfap-53* promoter sequence was fused the *wfap-53* coding sequence and GFP reporter. For microinjection, plasmid was purified using the QIAquick PCR Purification Kit (Qiagen, #28104) and co-injected with *rol-6* (su1006) into the gonads of young adult *C. elegans* hermaphrodites. To establish differential WFAP-53 expression levels, the *Pwfap-53::wfap-53::GFP* plasmid was microinjected at 50 ng/μL for high expression or 5 ng/μL for low expression.

### Protein Expression, and Purification

*C. reinhardtii* FAP53 cDNA was cloned into pET.M.3C and expressed in *E. coli* (DE3). Basically, transformed bacteria were cultured at 37°C, 200 rpm and induced by 0.5 mM IPTG at 25°C, 150 rpm overnight. Bacteria were lysed in lysis buffer (50 mM HEPES pH 7.5, 500 mM KCl, 10 mM imidazole, 5 mM β-ME, 100 μM PMSF and protease inhibitor (Roch)) and applied to Ni-NTA columns for purification. After washing by wash buffer (50 mM HEPES pH 7.5, 250 mM KCl, 40 mM imidazole, 5 mM β-ME), proteins were eluted by elution buffer (50 mM HEPES pH 7.5, 100 mM KCl, 200 mM imidazole, 5 mM β-ME). Eluted proteins were applied to a 10 kDa centricon to remove the imidazole. Protein concentration was determined by standard Bradford assay.

### Microtubule assembly assay

Tubulin (5 mg/mL) were supplemented with GTP (1 mM), MgCl_2_ (4 mM) and DMSO (4%) and incubated for 5 minutes on ice and then polymerized at 37°C for 30 minutes. Polymerized microtubules were supplemented with taxol (20 μM final) and centrifuged at 13,000 g with a tabletop centrifuge for 15 minutes at 37°C to remove unpolymerized tubulin. The pellet was further resuspended in taxol (20 μM final) supplemented BRB80. For microtubule doublet assembly, polymerized microtubules were mixed equivalently with unpolymerized tubulin (5 mg/mL) and add 1/138 molecular ratio of purified FAP53. The mixture was supplemented with GTP (1 mM), MgCl_2_ (4 mM) and DMSO (4%) incubated at room temperature for 5 minutes and then polymerized at 37°C for 30 minutes. Unpolymerized tubulin were removed by centrifugation.

### Live-Cell Imaging

Young adult *C. elegans* hermaphrodites were anesthetized with 0.1 mmol/L levamisole in M9 buffer, mounted on 3% agar pads. Images were captured using a spinning disk confocal system, which contains an Axio Observer Z1 microscope (Carl Zeiss) equipped with a 100×, 1.46 NA objective lens, an EMCCD camera (iXon+DU-897D-C00-#BV-500; Andor Technology), and the 405-nm, 488-nm and 568-nm lines of a Sapphire CW CDRH USB Laser System (Coherent) with a spinning disk confocal scan head (CSU-X1 Spinning Disk Unit; Yokogawa Electric Corporation). The confocal system was controlled by μManager (https://www.micro-manager.org), and time-lapse images were obtained using a 200 ms exposure time. Images were processed and analyzed with the ImageJ software (http://rsbweb.nih.gov/ij/).

### Cell Culture and Transfection

HeLa cells were cultured at 37 °C in a humidified incubator with 5% CO₂. Cells were maintained in Dulbecco’s Modified Eagle Medium (DMEM; Thermo Fisher Scientific) supplemented with 10% fetal bovine serum (FBS) and 1% penicillin–streptomycin (PS). For transient transfection, cells were seeded onto glass-bottom dishes and transfected with 500 ng of plasmid DNA using Lipofectamine 3000 (Thermo Fisher Scientific, Cat# L3000015) according to the manufacturer’s instructions. Cells were incubated for 16 hours post-transfection prior to fixation or live imaging.

### Fluorescence Microscopy

Confocal fluorescence imaging was performed using a Zeiss LSM 900 laser scanning confocal microscope equipped with Airyscan and a Plan-Apochromat 63×/1.4 NA oil immersion objective, operated with ZEN software (Carl Zeiss). EGFP and mScarlet were excited using 488 nm and 561 nm lasers, respectively, and Tubulin Tracker Deep Red (Thermo Fisher Scientific) was excited with a 640 nm laser. All images within the same experimental set were acquired under identical acquisition settings, including laser power, detector gain, and exposure time. Post-acquisition processing and image adjustments (e.g., brightness and contrast) were performed uniformly across samples using Fiji (ImageJ).

### Cryo-electron microscopy and tomography

Samples for cryo-EM were prepared using Vitrobot System (Thermo Scientific). Briefly, polymerized microtubule samples (5 μL) were dropped onto a glow-discharged cryo-EM grid (200 mesh, C-Flat Holey Carbon Grid Gold, EMS) and allowed to deposit for 1 minute. Then followed the standard operations of the system with setting the humidity to 100%, the blot time to 4 seconds, and the blot force to 2.

Cryo-electron microscopy images (Fig. 2B) were acquired with a Tecnai F20 TEM D545 (Thermo Scientific) cooled at the temperature of the liquid nitrogen and operating at 200 kV. Cryo-electron tomograms (Fig. 2C) were acquired with a 300 KV Titan Krios Microscopy (Thermo Scientific) equipped with a Cs corrector, GIF Quantum energy filter (Gatan Inc.), and K2 Summit direct electron detector (Gatan Inc.). Tilt series were collected from -60° to +60° at 33,000x of magnification (pixel size = 3.42 Å) with a defocus range of -2 to -4 µm for a total dose of 20 e-/Å2. The cryo-electron tomograms were aligned using patch tracking and reconstructed by R-weighted back projection using IMOD ^37^.

### TEM of chemical-fixed microtubules

Polymerized microtubule samples (5 μL) were dropped onto a glass-bottomed petri dish and allowed to deposit for 1 minute. Excess liquid was blotted, followed by immersion in 1 mL 2.5% (v/v) glutaraldehyde (Electron Microscopy Sciences) solution supplemented with 0.1% (w/v) tannic acid (Sigma) in PB buffer for 5 minutes. The fixative was then replaced with fresh 2.5% (v/v) glutaraldehyde in PB buffer, and fixed for an additional 1 hour at room temperature. After fixation, samples were washed three times with PB buffer, and were osmicated with 1% osmium tetroxide/1.5% potassium ferricyanide (Electron Microscopy Sciences) in distilled water for 30 minutes. Subsequent dehydration was performed using a graded ethanol series (50%, 70%, 90%, and 100%, 10 minutes per concentration). For resin embedding, samples were infiltrated with SPI-Pon 812 Resin (Structure Probe, Inc.) through a diluted series of ethanol/SPI-Pon 812 at 1:1, 1:2 and 1:3 ratio for 1 hour each, and then in pure resin overnight. After that, pure resin was changed once in the first 1 hour and then polymerize at 60°C for 48 hours. 70-nm ultrathin sections were cut by UC7 ultramicrotome (Leica Microsystems) and stained by 2% uranyl acetate and Reynold’s lead citrate. Sections were imaged on FEI Tecnai G2 Spirit (120 kV) electron microscope (Thermo Fisher Scientific Co.).

### TEM of C. elegans

TEM experiment was performed as an early protocol ^38^. Briefly, adult worms were loaded onto a 50 μm deep specimen carrier and rapid frozen with HPM100 high-pressure freezing (HPF) machine (Leica Microsystems). The specimens were then transferred into cryovials containing 1 mL 1% osmium tetroxide/0.1% uranyl acetate (Electron Microscopy Sciences) and processed in an AFS2 machine (Leica Microsystems) using a standard substitution and fixation program: -90°C for 48 hours, -60°C for 24 hours, -30°C for 18 hours, and finally rise to 4°C. Fixed specimens were washed three times with pure acetone, infiltrated with SPI-Pon 812 Resin (Structure Probe Inc.), embedded into a flat mold, and polymerized at 60°C for 3 days. 90-nm ultrathin sections were obtained using UC7 Ultramicrotome (Leica Microsystems) and picked onto 200 mesh copper grids (Gilder Grids Ltd.). Sections were poststained with 2% uranyl acetate and Reynold’s lead citrate and imaged on FEI Tecnai G2 Spirit (120 kV) electron microscope (Thermo Fisher Scientific Co.).

### Molecular dynamics simulations

Molecular models of GTP-tubulin dimer docking onto the A10-A11 protofilament patch was based on 6U42 PDB structures ^16^. The structures of TBB-4 and TBA-5 tubulin binding to Mg^2+^-GTP predicted by AlphaFold 3 (www.alphafoldserver.com) ^39^ were used as alpha- and beta-tubulin. The tubulins were then aligned to the outer junction of 6U42 model. Protonation state of ionizable amino acid residues were predicted by PDBFixer ^40^. To simplify the model, the ligands on A10-A11 protofilament were removed. The modeling of FAP53 residues 2-40 involved structural alignment with B01 tubulin dimer termini (plus/minus end), guided by their 6U42 conformation.

A virtual three-dimensional cubic reaction volume filled with TIP3P water with periodic boundary conditions was used for the simulation. The size of the reaction volume was set in such a way that the distance from the protein surface to the nearest box boundary was not initially less than two nanometers. The ionic strength of the solution was set at 100 mM by adding K^+^ and Cl^-^ ions and the total charge of the system was zero. Simulations were performed using the GROMACS 2024.2 software package with the CHARMM36m force field ^41^. The parameters of the GTP were generated using CHARMM General Force Field (CGenFF) by CHARMM-GUI ^42,43^.

After preparing each of the tubulin systems as described above, steepest descent algorithm was used to minimize the energy of the whole system. First, 100 ps long simulations were conducted with V-rescale thermostat (τ = 0.1 ps, T = 300 K). Second, we carried out another 100 ps simulations with the same V-rescale thermostat and C-rescale barostat (τ = 2.0 ps, compressibility = 4.5×10^-5^ bar^-1^, pressure = 1 bar). Position restrain was applied on protein heavy atoms during the two steps above. The production simulation runs were carried out in NPT ensemble at 300K, using V-rescale thermostat and C-rescale barostat for a duration of 100 ns each. In the simulations, positional restraints were applied to all Cα atoms of A10-A11 complex, except for the tubulin tails (refer to Figure 1C) of the tubulins in the A10-A11 protofilament patch, to mimic the stable state of the A-tubule as a microtubule wall fragment. Positional restraints were also applied from residue 34 to 40 in FAP53 protein to mimic the stable inner-microtubule interactions between FAP53 and other microtubule inner proteins. No positional restraints were applied to B01 dimer so that the interaction between B01 and A10-A11 could be observed. The Particle Mesh Ewald method was used to treat the long-range electrostatics. H-bond LINCS constraints allowed MD with 2 fs time step. PyMOL (version 2.0 Schrödinger, LLC) were used for visualization.

### Free energy landscape estimation

A Cartesian three-dimensional coordinate system was established: the Z-axis was oriented along the microtubule, pointing toward its plus end. The X-axis lay tangential to the microtubule, while the Y-axis was perpendicular to the X-axis. The origin of the coordinate system was defined as the initial center of mass of B01. During the 100 ns simulation, the center of mass coordinates of B01 was projected to the X-Y plane. The probability density was estimated by Kernel Density Estimation with Gaussian kernel. Bandwidth was determined. The probability density of (x, y) coordinate on the X-Y plane can be estimated as:

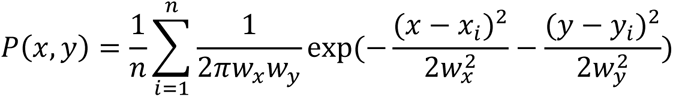

Here the bandwidth w is determined by Scott’s method:

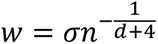

and the free energy was inferred from the probability density by Boltzmann inversion:

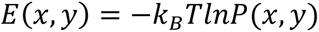

### Contact surface area calculation

The total contact area between B01 and the A-tubule was calculated by estimating the Solvent Accessible Surface Area (SASA) of each component in the complex. We define *A*_*a*−*b*_ as the contact area between components *a* and *b*, and *S*_*a*_ as the SASA of component *a*. The total contact area was computed using the following equation:

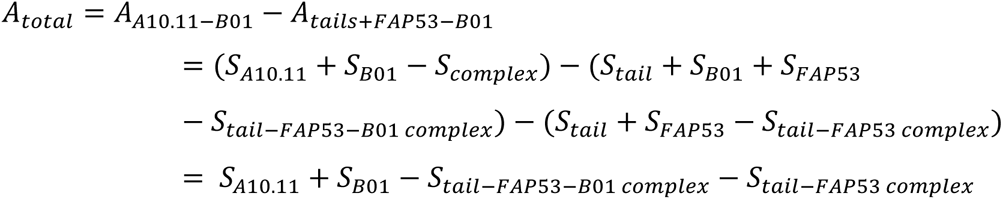

SASA values were computed using the GROMACS SASA module with default parameters.

## Acknowledgments

We thank the Tsinghua University Cryo-EM Facility of China National Center for Protein Sciences (Beijing) for HPF and EM data collection. This work was supported by the National Key R&D Program of China Grants 2024YFA1307301, 2022YFA1302700, 2019YFA0508401; and National Natural Science Foundation of China Grants 92254306, 31991190, 32270773, 32470730, 32070706 32270721, 32430026, and 32400610; and Pillars of the Nation Funding for Life Sciences, and TsienTang Life Science Development Fund at Tsinghua University.

---

**Figure S1.**
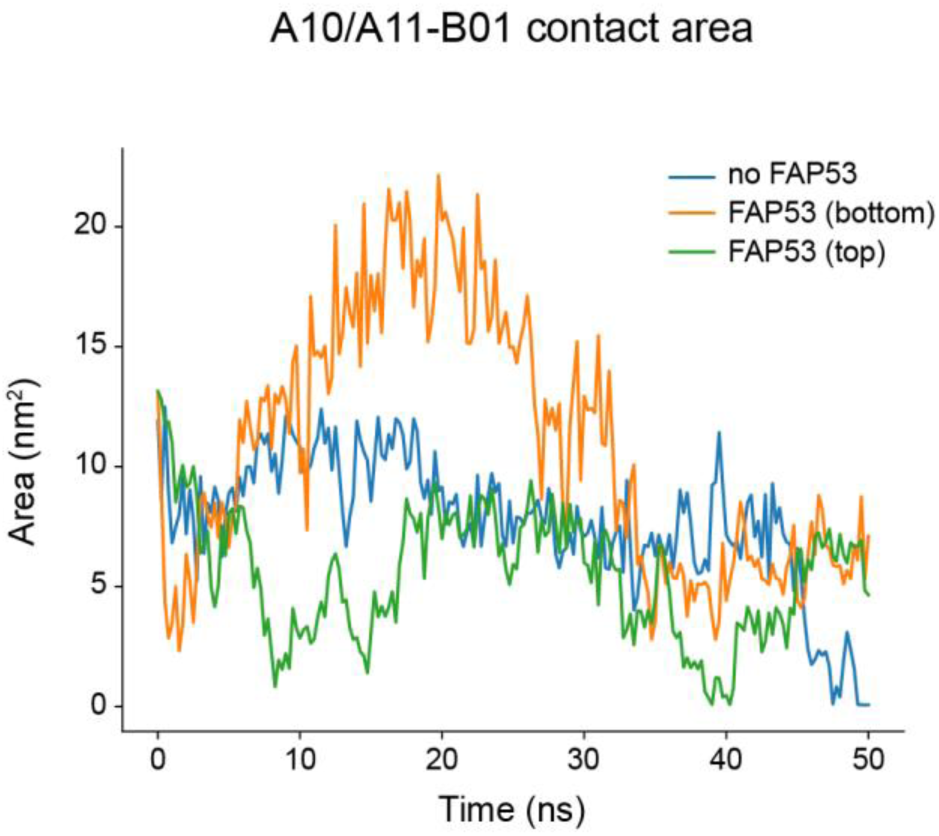
Molecular dynamic simulation predicts that FAP53 mitigates the inhibitory effect of CTT on formation of MTs assembly. The contact areas between B01 dimer and A10-A11 patch in different cases were showed. The first 50 ns of the simulations were discarded as the equilibration phase. The contact area was calculated every 0.5 ns during the last 50 ns.

**Figure S2.**
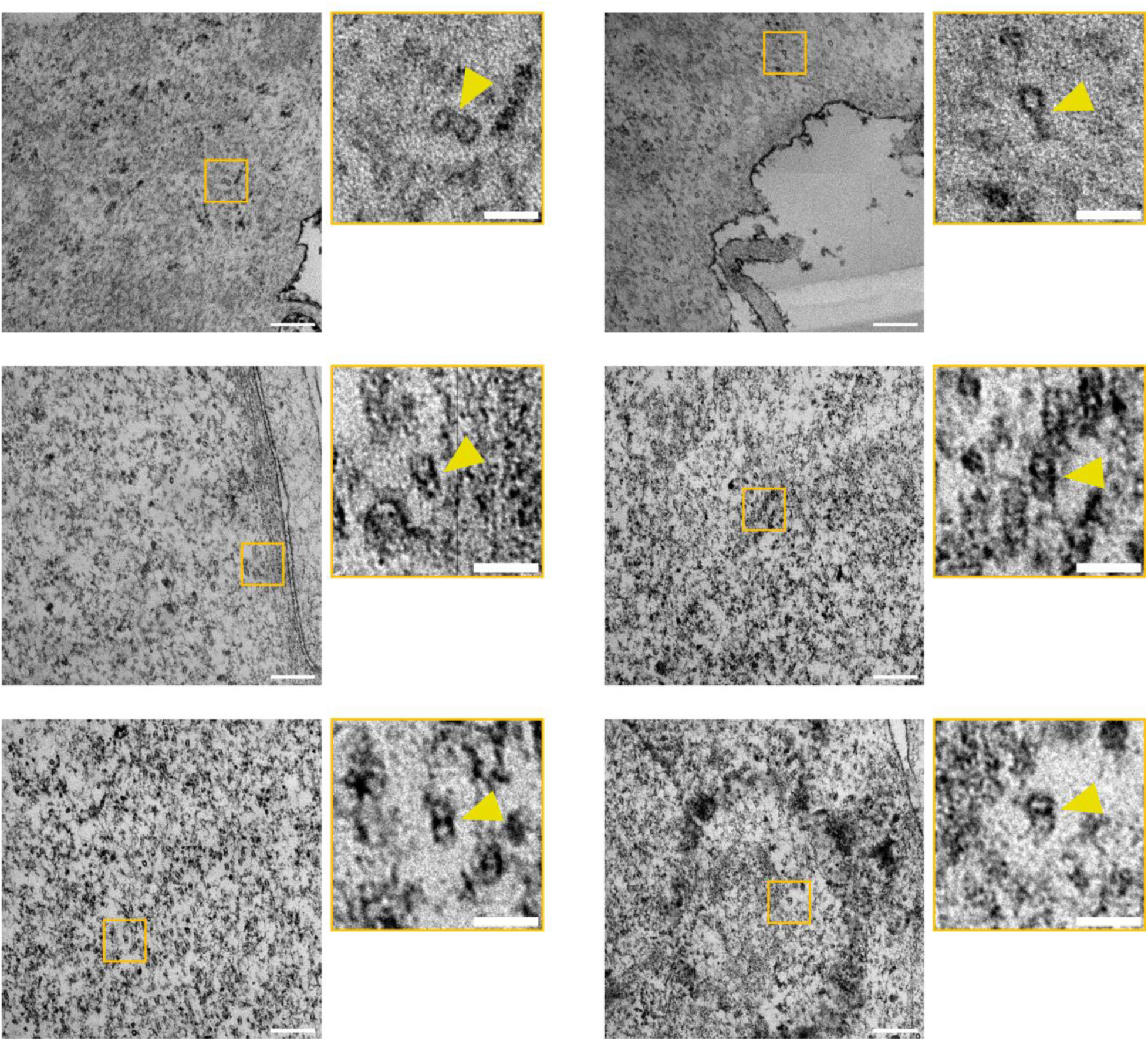
TEM images (cross sections) of the CFAP53+CFAP20 overexpression cells. High magnification on the right side shows the MTDs (arrowheads). Scale bars: 200 nm (low magnification), 50 nm (high magnification).

**Figure S3.**
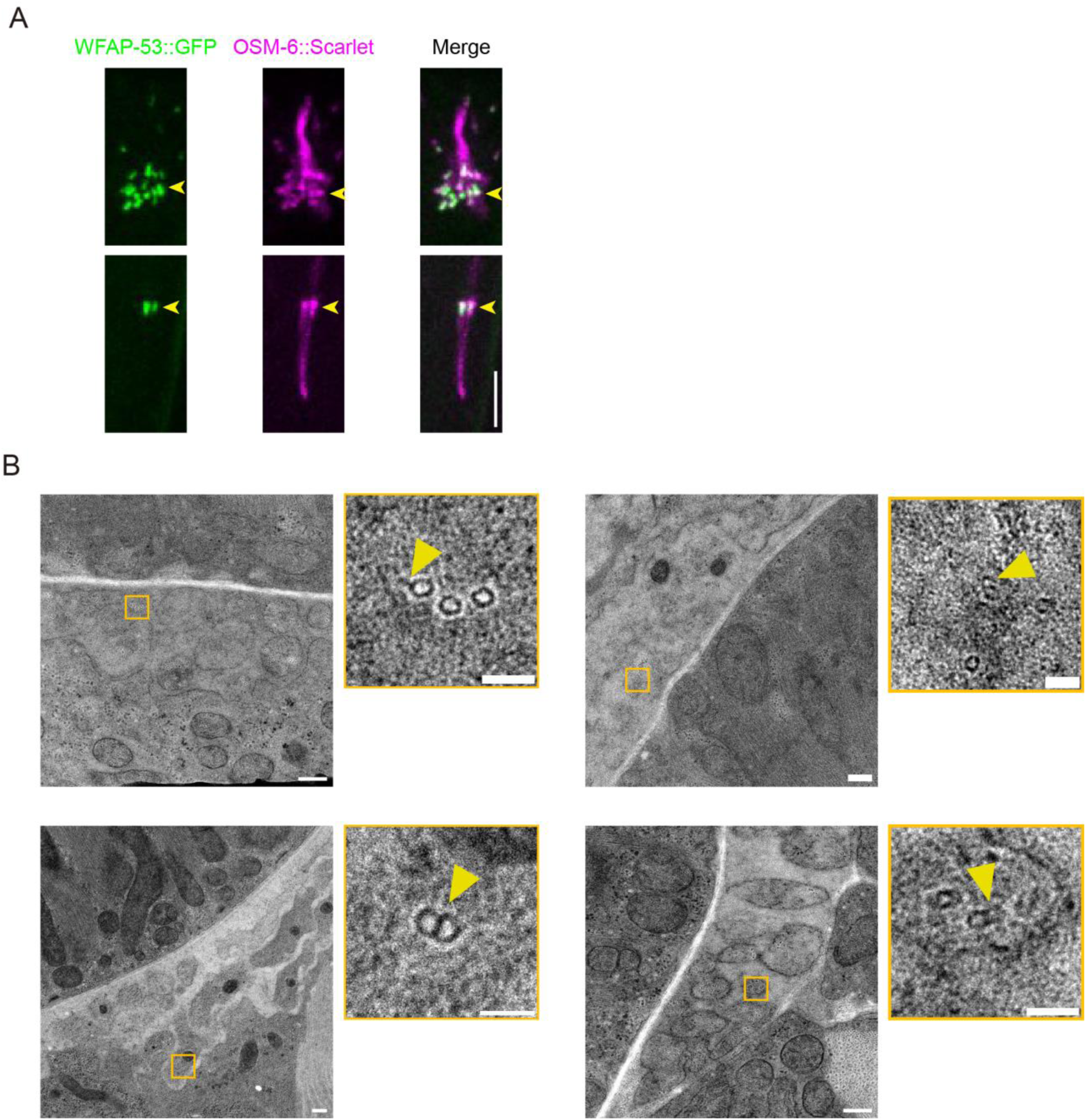
Overexpression of FAP53 structural homolog *wfap-53* induces ectopic MTDs in *C. elegans*. (A) Localization of WFAP-53 in amphid and phasmid cilia. Scale bar: 5 μm. (B) TEM images (cross sections) of neuronal dendrites in *wfap-53* overexpression animals. High magnification on the right side shows the ectopic MTDs. Scale bars: 200 nm (low magnification), 50 nm (high magnification).

**Table S1.** Strains and plasmids in this study.

| *C. elegans* strains used in this study |  |  |  |
| --- | --- | --- | --- |
| Strain name | Genotype | Method | Resource |
| GOU6769 | *casEx7016(Pwfap-53::wfap-53::GFP (50 ng/μL);rol-6(su1006)(+))* | Microinjection | This study |
| GOU6770 | *casEx7017(Pwfap-53::wfap-53::GFP (5 ng/μL);rol-6(su1006)(+))* | Microinjection | This study |

**Video S1.** Molecular dynamics simulation of the no FAP53 model. Top view (left) and side view (right) are showed. Duration: 100 ns. Related to Figure 1C.

**Video S2.** Molecular dynamics simulation of the FAP53 (bottom) model. Top view (left) and side view (right) are showed. Duration: 100 ns. Related to Figure 1C.

**Video S3.** Molecular dynamics simulation of the FAP53 (top) model. Top view (left) and side view (right) are showed. Duration: 100 ns. Related to Figure 1C.

## Reference

1. Hilgendorf, K.I., Myers, B.R., and Reiter, J.F. (2024). Emerging mechanistic understanding of cilia function in cellular signalling. Nat Rev Mol Cell Biol 25, 555–573. 10.1038/s41580-023-00698-5.

2. Klena, N., and Pigino, G. (2022). Structural Biology of Cilia and Intraflagellar Transport. Annu Rev Cell Dev Biol 38, 103–123. 10.1146/annurev-cellbio-120219-034238.

3. Lacey, S.E., and Pigino, G. (2025). The intraflagellar transport cycle. Nat Rev Mol Cell Biol 26, 175–192. 10.1038/s41580-024-00797-x.

4. Mill, P., Christensen, S.T., and Pedersen, L.B. (2023). Primary cilia as dynamic and diverse signalling hubs in development and disease. Nat Rev Genet 24, 421–441. 10.1038/s41576-023-00587-9.

5. Ou, G., and Scholey, J.M. (2022). Motor Cooperation During Mitosis and Ciliogenesis. Annu Rev Cell Dev Biol 38, 49–74. 10.1146/annurev-cellbio-121420-100107.

6. Rosenbaum, J.L., Moulder, J.E., and Ringo, D.L. (1969). Flagellar elongation and shortening in Chlamydomonas. The use of cycloheximide and colchicine to study the synthesis and assembly of flagellar proteins. J Cell Biol 41, 600–619. 10.1083/jcb.41.2.600.

7. Rao, V.G., Subramanianbalachandar, V.A., Magaj, M.M., Redemann, S., and Kulkarni, S.S. (2025). Mechanisms of cilia regeneration in Xenopus multiciliated epithelium in vivo. EMBO Rep 26, 2192–2220. 10.1038/s44319-025-00414-8.

8. Huang, B., Rifkin, M.R., and Luck, D.J. (1977). Temperature-sensitive mutations affecting flagellar assembly and function in Chlamydomonas reinhardtii. J Cell Biol 72, 67–85. 10.1083/jcb.72.1.67.

9. Li, M., Chen, Z., Guo, Z., Wang, Y., Chai, Y., Li, W., and Ou, G. (2025). Alpha-tubulin tails regulate axoneme differentiation. Proc Natl Acad Sci U S A 122, e2414731122. 10.1073/pnas.2414731122.

10. Schmidt-Cernohorska, M., Zhernov, I., Steib, E., Le Guennec, M., Achek, R., Borgers, S., Demurtas, D., Mouawad, L., Lansky, Z., Hamel, V., and Guichard, P. (2019). Flagellar microtubule doublet assembly in vitro reveals a regulatory role of tubulin C-terminal tails. Science 363, 285–288. 10.1126/science.aav2567.

11. Linck, R.W., and Langevin, G.L. (1981). Reassembly of flagellar B (alpha beta) tubulin into singlet microtubules: consequences for cytoplasmic microtubule structure and assembly. J Cell Biol 89, 323–337. 10.1083/jcb.89.2.323.

12. Serrano, L., de la Torre, J., Maccioni, R.B., and Avila, J. (1984). Involvement of the carboxyl-terminal domain of tubulin in the regulation of its assembly. Proc Natl Acad Sci U S A 81, 5989–5993. 10.1073/pnas.81.19.5989.

13. Gui, M., Farley, H., Anujan, P., Anderson, J.R., Maxwell, D.W., Whitchurch, J.B., Botsch, J.J., Qiu, T., Meleppattu, S., Singh, S.K., et al. (2021). De novo identification of mammalian ciliary motility proteins using cryo-EM. Cell 184, 5791–5806 e5719. 10.1016/j.cell.2021.10.007.

14. Kubo, S., Black, C.S., Joachimiak, E., Yang, S.K., Legal, T., Peri, K., Khalifa, A.A.Z., Ghanaeian, A., McCafferty, C.L., Valente-Paterno, M., et al. (2023). Native doublet microtubules from Tetrahymena thermophila reveal the importance of outer junction proteins. Nat Commun 14, 2168. 10.1038/s41467-023-37868-0.

15. Leung, M.R., Zeng, J., Wang, X., Roelofs, M.C., Huang, W., Zenezini Chiozzi, R., Hevler, J.F., Heck, A.J.R., Dutcher, S.K., Brown, A., et al. (2023). Structural specializations of the sperm tail. Cell 186, 2880–2896 e2817. 10.1016/j.cell.2023.05.026.

16. Ma, M., Stoyanova, M., Rademacher, G., Dutcher, S.K., Brown, A., and Zhang, R. (2019). Structure of the Decorated Ciliary Doublet Microtubule. Cell 179, 909–922 e912. 10.1016/j.cell.2019.09.030.

17. Walton, T., Gui, M., Velkova, S., Fassad, M.R., Hirst, R.A., Haarman, E., O’Callaghan, C., Bottier, M., Burgoyne, T., Mitchison, H.M., and Brown, A. (2023). Axonemal structures reveal mechanoregulatory and disease mechanisms. Nature 618, 625–633. 10.1038/s41586-023-06140-2.

18. Zhou, L., Liu, H., Liu, S., Yang, X., Dong, Y., Pan, Y., Xiao, Z., Zheng, B., Sun, Y., Huang, P., et al. (2023). Structures of sperm flagellar doublet microtubules expand the genetic spectrum of male infertility. Cell 186, 2897–2910 e2819. 10.1016/j.cell.2023.05.009.

19. Ide, T., Twan, W.K., Lu, H., Ikawa, Y., Lim, L.X., Henninger, N., Nishimura, H., Takaoka, K., Narasimhan, V., Yan, X., et al. (2020). CFAP53 regulates mammalian cilia-type motility patterns through differential localization and recruitment of axonemal dynein components. PLoS Genet 16, e1009232. 10.1371/journal.pgen.1009232.

20. Lu, H., Twan, W.K., Ikawa, Y., Khare, V., Mukherjee, I., Schou, K.B., Chua, K.X., Aqasha, A., Chakrabarti, S., Hamada, H., and Roy, S. (2024). Localisation and function of key axonemal microtubule inner proteins and dynein docking complex members reveal extensive diversity among vertebrate motile cilia. Development 151. 10.1242/dev.202737.

21. Narasimhan, V., Hjeij, R., Vij, S., Loges, N.T., Wallmeier, J., Koerner-Rettberg, C., Werner, C., Thamilselvam, S.K., Boey, A., Choksi, S.P., et al. (2015). Mutations in CCDC11, which encodes a coiled-coil containing ciliary protein, causes situs inversus due to dysmotility of monocilia in the left-right organizer. Hum Mutat 36, 307–318. 10.1002/humu.22738.

22. Silva, E., Betleja, E., John, E., Spear, P., Moresco, J.J., Zhang, S., Yates, J.R., 3rd, Mitchell, B.J., and Mahjoub, M.R. (2016). Ccdc11 is a novel centriolar satellite protein essential for ciliogenesis and establishment of left-right asymmetry. Mol Biol Cell 27, 48–63. 10.1091/mbc.E15-07-0474.

23. Guo, Z., Tan, M., Zhu, H., Lou, G., Xia, X., Yang, W., Lv, Y., Huang, J., Wang, R., Hao, B., and Liao, S. (2025). Identification of novel biallelic mutations in CFAP53 associated with fetal situs inversus totalis and literature review. J Appl Genet. 10.1007/s13353-025-00950-y.

24. Gur, M., Cohen, E.B., Genin, O., Fainsod, A., Perles, Z., and Cinnamon, Y. (2017). Roles of the cilium-associated gene CCDC11 in left-right patterning and in laterality disorders in humans. Int J Dev Biol 61, 267–276. 10.1387/ijdb.160442yc.

25. Noel, E.S., Momenah, T.S., Al-Dagriri, K., Al-Suwaid, A., Al-Shahrani, S., Jiang, H., Willekers, S., Oostveen, Y.Y., Chocron, S., Postma, A.V., et al. (2016). A Zebrafish Loss-of-Function Model for Human CFAP53 Mutations Reveals Its Specific Role in Laterality Organ Function. Hum Mutat 37, 194–200. 10.1002/humu.22928.

26. Perles, Z., Cinnamon, Y., Ta-Shma, A., Shaag, A., Einbinder, T., Rein, A.J., and Elpeleg, O. (2012). A human laterality disorder associated with recessive CCDC11 mutation. J Med Genet 49, 386–390. 10.1136/jmedgenet-2011-100457.

27. Satoda, Y., Noguchi, T., Fujii, T., Taniguchi, A., Katoh, Y., and Nakayama, K. (2022). BROMI/TBC1D32 together with CCRK/CDK20 and FAM149B1/JBTS36 contributes to intraflagellar transport turnaround involving ICK/CILK1. Mol Biol Cell 33, ar79. 10.1091/mbc.E22-03-0089.

28. Guo, Z., Wang, Y., and Ou, G. (2024). Utilizing the scale-invariant feature transform algorithm to align distance matrices facilitates systematic protein structure comparison. Bioinformatics 40. 10.1093/bioinformatics/btae064.

29. Taylor, S.R., Santpere, G., Weinreb, A., Barrett, A., Reilly, M.B., Xu, C., Varol, E., Oikonomou, P., Glenwinkel, L., McWhirter, R., et al. (2021). Molecular topography of an entire nervous system. Cell 184, 4329–4347 e4323. 10.1016/j.cell.2021.06.023.

30. Chen, Z., Li, M., Zhu, H., and Ou, G. (2023). Modulation of inner junction proteins contributes to axoneme differentiation. Proc Natl Acad Sci U S A 120, e2303955120. 10.1073/pnas.2303955120.

31. Dymek, E.E., Lin, J., Fu, G., Porter, M.E., Nicastro, D., and Smith, E.F. (2019). PACRG and FAP20 form the inner junction of axonemal doublet microtubules and regulate ciliary motility. Mol Biol Cell 30, 1805–1816. 10.1091/mbc.E19-01-0063.

32. Yanagisawa, H.A., Mathis, G., Oda, T., Hirono, M., Richey, E.A., Ishikawa, H., Marshall, W.F., Kikkawa, M., and Qin, H. (2014). FAP20 is an inner junction protein of doublet microtubules essential for both the planar asymmetrical waveform and stability of flagella in Chlamydomonas. Mol Biol Cell 25, 1472–1483. 10.1091/mbc.E13-08-0464.

33. Bangera, M., Dungdung, A., Prabhu, S., and Sirajuddin, M. (2023). Doublet microtubule inner junction protein FAP20 recruits tubulin to the microtubule lattice. Structure 31, 1535–1544 e1534. 10.1016/j.str.2023.09.010.

34. Owa, M., Uchihashi, T., Yanagisawa, H.A., Yamano, T., Iguchi, H., Fukuzawa, H., Wakabayashi, K.I., Ando, T., and Kikkawa, M. (2019). Inner lumen proteins stabilize doublet microtubules in cilia and flagella. Nat Commun 10, 1143. 10.1038/s41467-019-09051-x.

35. Reiter, J.F., and Leroux, M.R. (2017). Genes and molecular pathways underpinning ciliopathies. Nat Rev Mol Cell Biol 18, 533–547. 10.1038/nrm.2017.60.

36. Pazour, G.J. (2024). Cilia Structure and Function in Human Disease. Curr Opin Endocr Metab Res 34. 10.1016/j.coemr.2024.100509.

37. Kremer, J.R., Mastronarde, D.N., and McIntosh, J.R. (1996). Computer visualization of three-dimensional image data using IMOD. J Struct Biol 116, 71–76. 10.1006/jsbi.1996.0013.

38. Xie, C., Li, L., Li, M., Shao, W., Zuo, Q., Huang, X., Chen, R., Li, W., Brunnbauer, M., Okten, Z., et al. (2020). Optimal sidestepping of intraflagellar transport kinesins regulates structure and function of sensory cilia. EMBO J 39, e103955. 10.15252/embj.2019103955.

39. Abramson, J., Adler, J., Dunger, J., Evans, R., Green, T., Pritzel, A., Ronneberger, O., Willmore, L., Ballard, A.J., Bambrick, J., et al. (2024). Accurate structure prediction of biomolecular interactions with AlphaFold 3. Nature 630, 493–500. 10.1038/s41586-024-07487-w.

40. Eastman, P., Swails, J., Chodera, J.D., McGibbon, R.T., Zhao, Y., Beauchamp, K.A., Wang, L.P., Simmonett, A.C., Harrigan, M.P., Stern, C.D., et al. (2017). OpenMM 7: Rapid development of high performance algorithms for molecular dynamics. PLoS Comput Biol 13, e1005659. 10.1371/journal.pcbi.1005659.

41. Huang, J., Rauscher, S., Nawrocki, G., Ran, T., Feig, M., de Groot, B.L., Grubmuller, H., and MacKerell, A.D., Jr. (2017). CHARMM36m: an improved force field for folded and intrinsically disordered proteins. Nat Methods 14, 71–73. 10.1038/nmeth.4067.

42. Jo, S., Kim, T., Iyer, V.G., and Im, W. (2008). CHARMM-GUI: a web-based graphical user interface for CHARMM. J Comput Chem 29, 1859–1865. 10.1002/jcc.20945.

43. Kim, S., Lee, J., Jo, S., Brooks, C.L., 3rd, Lee, H.S., and Im, W. (2017). CHARMM-GUI ligand reader and modeler for CHARMM force field generation of small molecules. J Comput Chem 38, 1879–1886. 10.1002/jcc.24829.

